# Effect of polyethylene glycol on growth of *Escherichia coli* DH5α and *Bacillus subtilis* NRS-762

**DOI:** 10.1101/2020.09.03.282376

**Authors:** Wenfa Ng

**Affiliations:** Department of Chemical and Biomolecular Engineering, National University of Singapore

**Keywords:** dose response, polyethylene glycol, toxicity, biomass formation, metabolism, optical density, pH variation, culture broth, *Escherichia coli*, *Bacillus subtilis*

## Abstract

Polyethylene glycol is commonly used in fermentation as an anti-foam for preventing the rise of foam to the top plate of the bioreactor, which increases contamination risk. However, its potential toxicity to growth of various microorganisms is not well understood at the species and strain level. Hence, the objective of this study was to understand the impact of different concentrations of polyethylene glycol at the 1, 5 and 10 g/L level on the aerobic growth of *Escherichia coli* DH5α and *Bacillus subtilis* NRS-762 in LB Lennox medium in shake flasks. Experiment results revealed that polyethylene glycol (PEG) (molecular weight ∼8000 Da), at all concentrations tested, did not affect biomass formation and metabolism in *E. coli* DH5α at 37 °C. This came about through the observation of similar maximal optical density obtained during growth of *E. coli* DH5α under differing concentrations of PEG. Furthermore, the anti-foam did not affect the pH profile. On the other hand, PEG did exhibit some toxicity towards the growth of *B. subtilis* NRS-762 in LB Lennox medium. Specifically, maximal optical density obtained decline with higher exposure to PEG in a concentration dependent manner, up to a threshold concentration of 5 g/L. For example, maximal optical density obtained in *B. subtilis* NRS-762 without addition of PEG was 4.4, but the value obtained with 1 g/L of the anti-foam decreased to 4.1 and a further 3.8 on exposure to 5 g/L and 10 g/L PEG. pH variation in culture broth, however, told a different story, where the profiles for exposure to PEG at all concentrations coincide with each other and was similar to the one without exposure to the anti-foam; thereby, suggesting that metabolic processes in *B. subtilis* NRS-762 were not significantly affected by exposure to PEG. Collectively, PEG anti-foam exerted species-specific toxicity effect on biomass formation, and possibly metabolism. The latter may not be sufficiently significant to affect the types of metabolites secreted by the bacterium, and thus, be detected by measurement of pH of culture broth. *E. coli* DH5α was better able to cope with PEG at all concentrations compared to *B. subtilis* NRS-762, which showed dose-dependent toxicity effect on biomass formation.

**Highlights:**

- Polyethylene glycol (molecular weight ∼ 8000 Da) did not affect aerobic growth of *Escherichia coli* DH5α at 37 °C in LB medium at all concentrations tested: 0, 1, 5, 10 g/L.
- Growth curves of the bacterium at different concentrations of polyethylene glycol (PEG) coincided with each other.
- Similar pH profiles were also obtained for *E. coli* DH5α growth in LB medium with different PEG concentrations.
- However, PEG exerted toxicity effect on *Bacillus subtilis* NRS-762 during growth in LB medium at 30 °C, with reduction of biomass formation in a dose dependent manner.
- Similar to the case for *E. coli* DH5α. pH profiles of *B. subtilis* NRS-762 coincided with each other irrespective of the concentrations of PEG used.

## Introduction

Polyethylene glycol (PEG) is commonly used as an anti-foam in microbial fermentation, especially at the bioreactor level, where profusion of dissolved oxygen and impellar stirring combined to result in rising of foam to the top plate of the bioreactor which increases the risk of contamination. To be useful as an anti-foam, PEG should not be biodegraded, but there have been reports of isolation of bacterial consortia capable of degrading PEG through action of PEG dehydrogenase.^1^

Known to be benign, PEG is approved as part of pharmaceutical formulations,^2^ but there have been emerging reports of its toxicity to cells. Reports in the literature suggests that derivatives of polyethylene glycol are cytotoxic to mammalian cells.^3,4^ Specifically, low molecular weight PEG has been shown to induce chromosomal aberrations in *in vitro* cultured Chinese Hamster epithelial liver cells.^4^ Furthermore, antibacterial effects of dental impacts coated with long-chain PEG have also been reported.^5^ This effect has been corroborated by other studies using PEG coated silver nanoparticles for killing multi-drug resistant bacterial species.^6^ Efficient killing of *Escherichia coli* DH5α and *Bacillus subtilis* suggests antibacterial properties of the PEG functionalized silver nanoparticles, but proven toxicity effect of silver on bacterial cells meant that effect of PEG coating on bacteria killing may be ambivalent.

Hence, there is a need to understand the effect of PEG on the growth of common bacteria such as *E. coli* and *B. subtilis*, which are commonly used in biotechnology for expressing recombinant proteins in bioreactor-based production of useful pharmaceuticals or value-added products. Such growth-related assays in liquid culture format would help disentangle the observed convoluted effect of antibacterial effects of PEG functionalized materials through the sole use of PEG.

Thus, using *Escherichia coli* DH5α (ATCC 53868) and *Bacillus subtilis* NRS-762 (ATCC 8473) as model organisms, this study aims to understand the effect of different concentrations of polyethylene glycol (0, 1, 5, and 10 g/L, average molecular weight 8000 Da) on the aerobic growth of *E. coli* DH5α and *B. subtilis* NRS-762 in LB Lennox medium contained in shake flasks. Specifically, optical density and pH measurement would be taken to ascertain the time course impact of exposure to different concentrations of polyethylene glycol on bacterial growth and metabolism. pH was used as a proxy parameter for the net amount of metabolites secreted into the culture broth during growth.

## Results and Discussion

Experiment results revealed that there was no difference in growth of *E. coli* DH5α with or without influence of PEG anti-foam (Figure 1). Specifically, growth curves of *E. coli* DH5α cultivated under 1, 5 and 10 g/L PEG coincided with each other, while that of *E. coli* DH5α cultured without the anti-foam yield the same maximal optical density of ∼3.3. Thus, PEG is non-toxic to *E. coli* DH5α, and its presence even at 10 g/L did not significantly disturb the osmotic balance of the culture medium for growth of *E. coli* DH5α.

**Figure 1:**
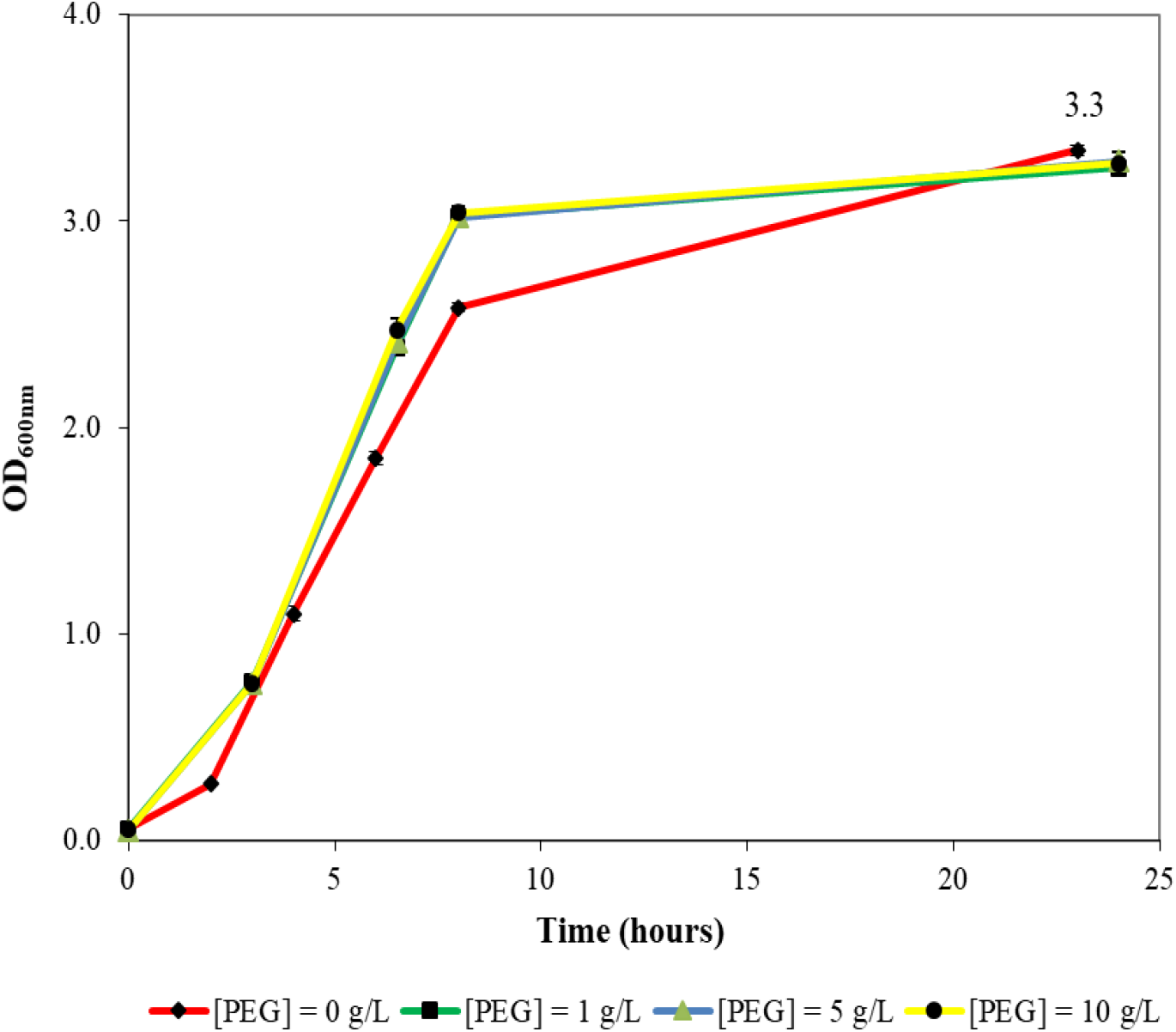
Growth of *E. coli* DH5α under varying concentrations of polyethylene glycol (PEG) in LB Lennox medium. There was essentially no difference to growth of *E. coli* DH5α with or without influence of the anti-foam, PEG.

Observation of pH variation during growth of *E. coli* DH5α in LB Lennox medium under varying concentrations of PEG revealed no difference to growth performance of the bacterium under different concentrations of the anti-foam (Figure 2). Specifically, the pH profile for growth under different PEG concentrations coincided with each other to a large degree; thus, highlighting that there were no significant changes in metabolism of cells during growth under different concentrations of PEG. In fact, the pH profile was the same when *E. coli* DH5α was grown without addition of PEG, which suggested that infusion of PEG did not affect the metabolism of the bacterium. However, small amounts of biofilm were found at the air-liquid interface of the shake flask at 8 hours of incubation under different concentrations of PEG. The amount of biofilm increased with time during culture under PEG influence.

**Figure 2:**
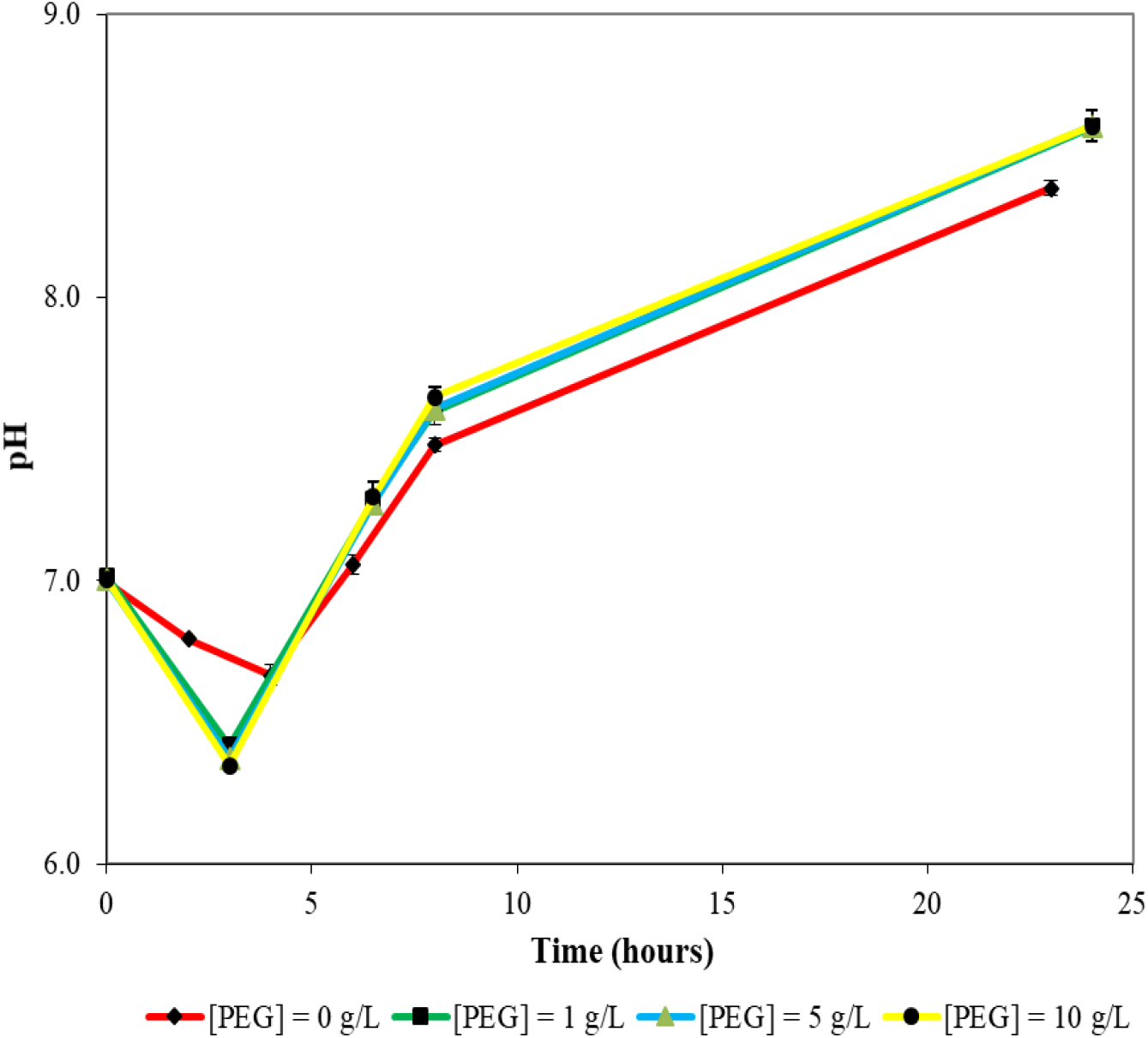
pH variation during culture of *E. coli* DH5α in LB Lennox medium at 37 °C under varying concentrations of PEG. Similar to the case for optical density, no difference in pH variation exists for culture of the bacterium under different concentrations of 1, 5 and 10 g/L PEG.

Growth of *B. subtilis* NRS-762 in LB Lennox medium under different concentrations of PEG revealed some toxicity effect of the anti-foam on growth of the bacterium (Figure 3). Specifically, without addition of PEG, the maximal optical density obtained was 4.4. This value decreased to 4.1 after addition of 1 g/L of PEG. Upon addition of 5 g/L and 10 g/L PEG, the maximal optical density attained decreased to 3.8, which highlighted that PEG exerted negative impacts to growth of *B. subtilis* NRS-762. This is in stark contrast to the case for *E. coli* DH5α. Besides reduction in maximal optical density obtained, growth was also slower with addition of PEG and the effect was concentration dependent, where higher concentration of PEG resulted in slower growth. However, a threshold existed at 5 g/L of PEG where there was no further concentration dependent effect. Specifically, both 5 g/L and 10 g/L PEG depressed growth rate and maximal optical density to the same extent.

**Figure 3:**
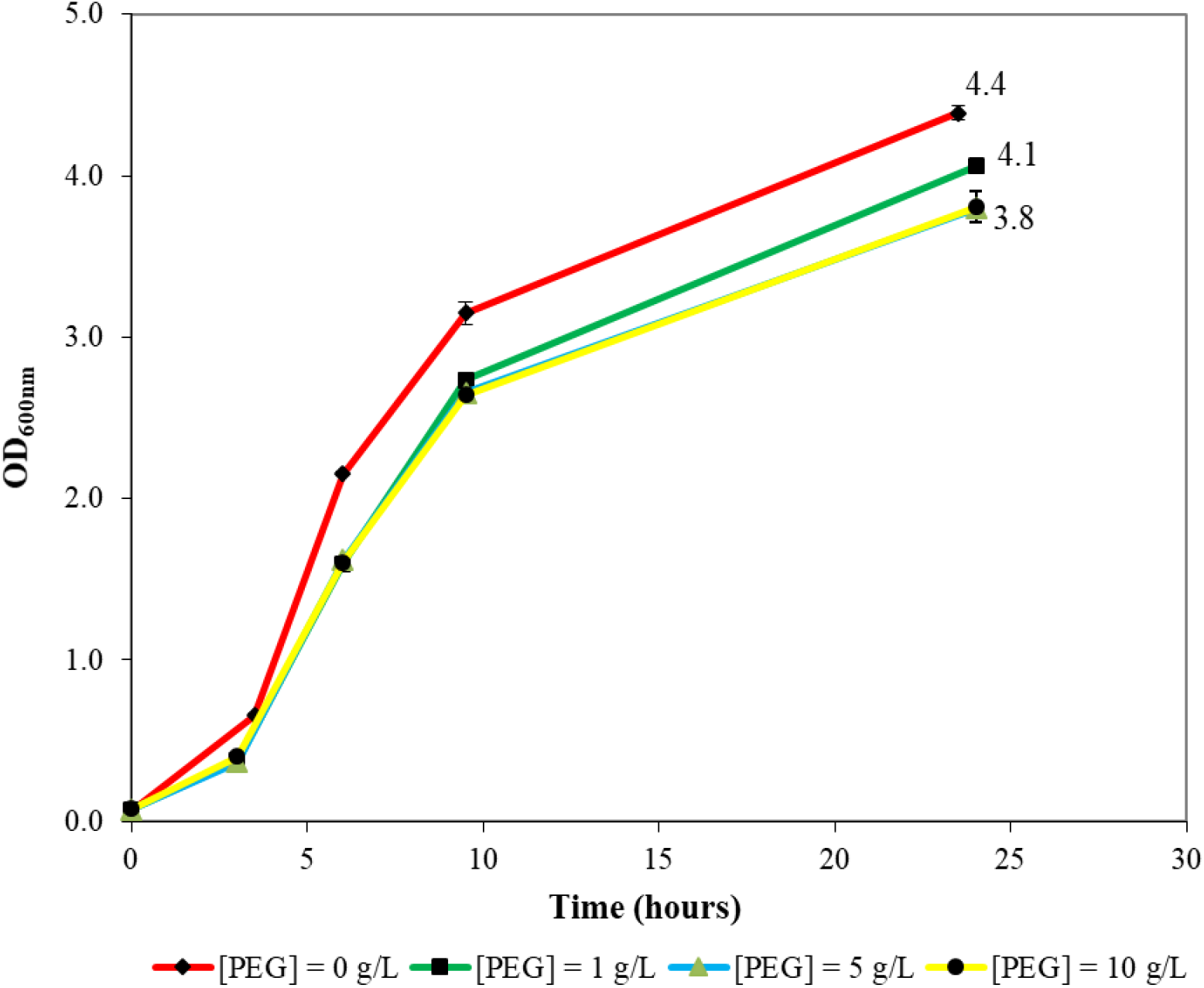
Aerobic growth of *B. subtilis* NRS-762 in LB Lennox medium under varying concentrations of PEG. Note that the maximal optical density obtained decreased in a concentration dependent manner with respect to PEG.

pH variation with culture time in cultures of *B. subtilis* NRS-762 exposed to different concentrations of PEG revealed that there was no concentration dependent impact on pH profile (Figure 4). Specifically, the same pH profile was obtained independent of the concentration of PEG added; thereby, highlighting that the anti-foam has minimal impact on the metabolism of the cells, except to reduce the biomass formation capacity of the bacterium under different PEG concentrations.

**Figure 4:**
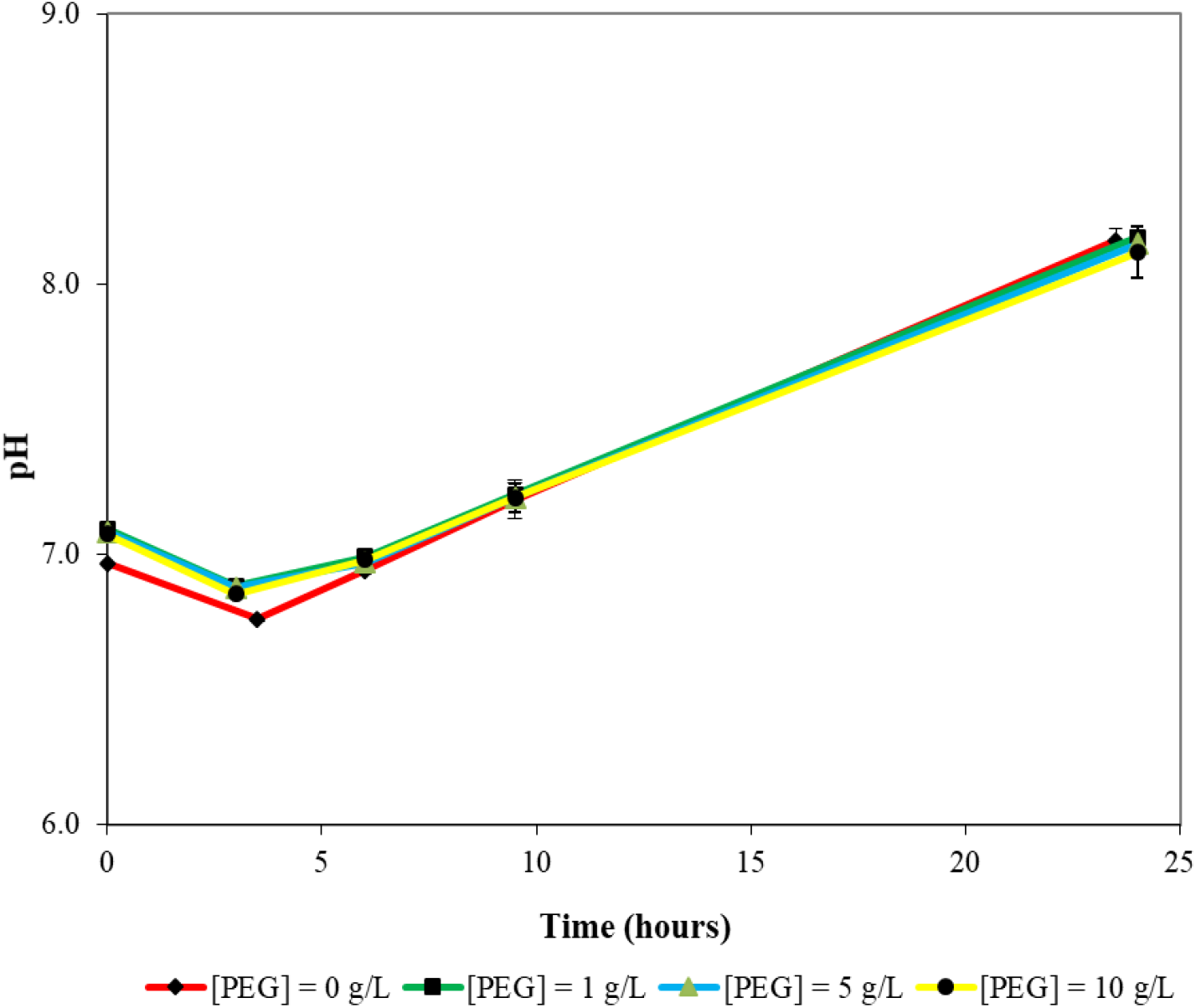
Variation of pH during growth of *B. subtilis* NRS-762 in LB Lennox medium at 30 °C under different concentrations of PEG. In general, the same trend holds independent of the concentration of PEG used; thereby, suggesting that the anti-foam has minimal impact on the metabolism of *B. subtilis* NRS-762.

## Conclusions

PEG did not exert significant negative impacts on growth of *E. coli* DH5α in LB Lennox medium at 37 °C, where there was no reduction of maximal optical density obtained and the pH variation with culture time indicated that the anti-foam did not affect the metabolism of the cells in a significant way. However, the anti-foam did reduce the maximal optical density of *B. subtilis* NRS-762 in a concentration dependent manner during aerobic growth of the bacterium in LB Lennox medium at 30 °C. Specifically, maximal optical density obtained decreased from 4.4 when there was no PEG added to 4.1 with 1 g/L of PEG added. Optical density of 3.8 was obtained when 5 g/L and 10 g/L of PEG was added into LB Lennox medium. However, toxicity impact of PEG did not manifest as changes to the pH profile obtained during growth of *B. subtilis* NRS-762 under differing concentrations of polyethylene glycol, which suggested that metabolism of cells were not significantly affected by the anti-foam. More importantly, decline in optical density with increasing exposure to PEG thus likely came from effect of the anti-foam on biomass formation processes.

## Materials and methods

### Materials

LB Lennox was purchased from Difco and used as is. Composition of the medium was in [g/L]: Tryptone, 10.0; Yeast extract, 5.0; NaCl, 5.0; Polyethylene glycol of molecular weight ∼8000 Da was purchased from Sigma-Aldrich. It was dissolved in deionized water and filter sterilized with a 0.22 µm Pall polyethersulfone membrane filter during addition to sterilized LB Lennox medium.

### Growth of bacteria in liquid medium

Stock cultures of *E. coli* DH5α and *B. subtilis* NRS-762 were prepared in glycerol and stored at – 70 °C prior to use. Glycerol stock cultures were used to inoculate seed cultures of 100 mL LB Lennox medium in 250 mL glass shake flask under aerobic culture conditions at 230 rpm rotational shaking in a temperature controlled incubator shaker. Incubation temperature was 37 °C for *E. coli* DH5α and 30 °C for *B. subtilis* NRS-762. After 24 hours of incubation, 1 mL of inoculum from the seed cultures was used in inoculating the experiment cultures. Three biological replicates were performed.

### Optical density and pH measurement

At appropriate time points, aliquots of culture broth were obtained for measuring optical density and pH. Optical density was measured using a Shimadzu Biospec Mini spectrophotometer with a quartz cuvette of pathlength of 10 mm. Appropriate dilution with deionized water was used if the optical density exceeded 1. pH was measured, without any sample dilution, using an Orion 9156 BNWP pH probe outfitted to a Mettler Toledo Delta 320 pH meter.

## Conflicts of interest

The author declares no conflicts of interest.

## Author’s contribution

The author designed and performed the experiments, analysed the data, and wrote the manuscript

## Funding

The author thank the National University of Singapore for financial support.

## Notes

### Competing Interest Statement

The authors have declared no competing interest.

